# CHARMM-GUI *Covalent Ligand Docker* as a Web-based Molecular Docking Platform for Covalent Ligands

**DOI:** 10.64898/2026.07.13.738313

**Authors:** Lingyang Kong, Donghyuk Suh, Wonpil Im

## Abstract

Covalent inhibitor research is an emerging topic in drug discovery due to its superior performance in specificity and inhibition effects. While molecular docking is a popular strategy in prediction and assessment of ligand conformations or poses in receptor proteins, covalent ligand docking requires nontrivial preparation efforts, as the ligand structure changes during the covalent complex formation. In order to facilitate molecular docking for covalent ligands, we have developed CHARMM-GUI *Covalent Ligand Docker* (CGUI-CLD), a new module for covalent ligand docking supported by AutoDock4. CGUI-CLD automates ligand preparation, supports ligand modification, implements docking simulation, and presents results through an intuitive user interface. A knowledge-based library built in CGUI-CLD currently supports 66 warheads and 8 amino acids, which can be used to automate the covalent ligand transformation from a pre-reaction to a post-reaction adduct form seamlessly. Moreover, CHARMM-GUI *High-Throughput Simulator* is integrated for rapid generation of multiple molecular dynamics simulation systems. CGUI-CLD is expected to significantly reduce a massive workload of covalent ligand docking and advance covalent ligand research.

Abstract TOC

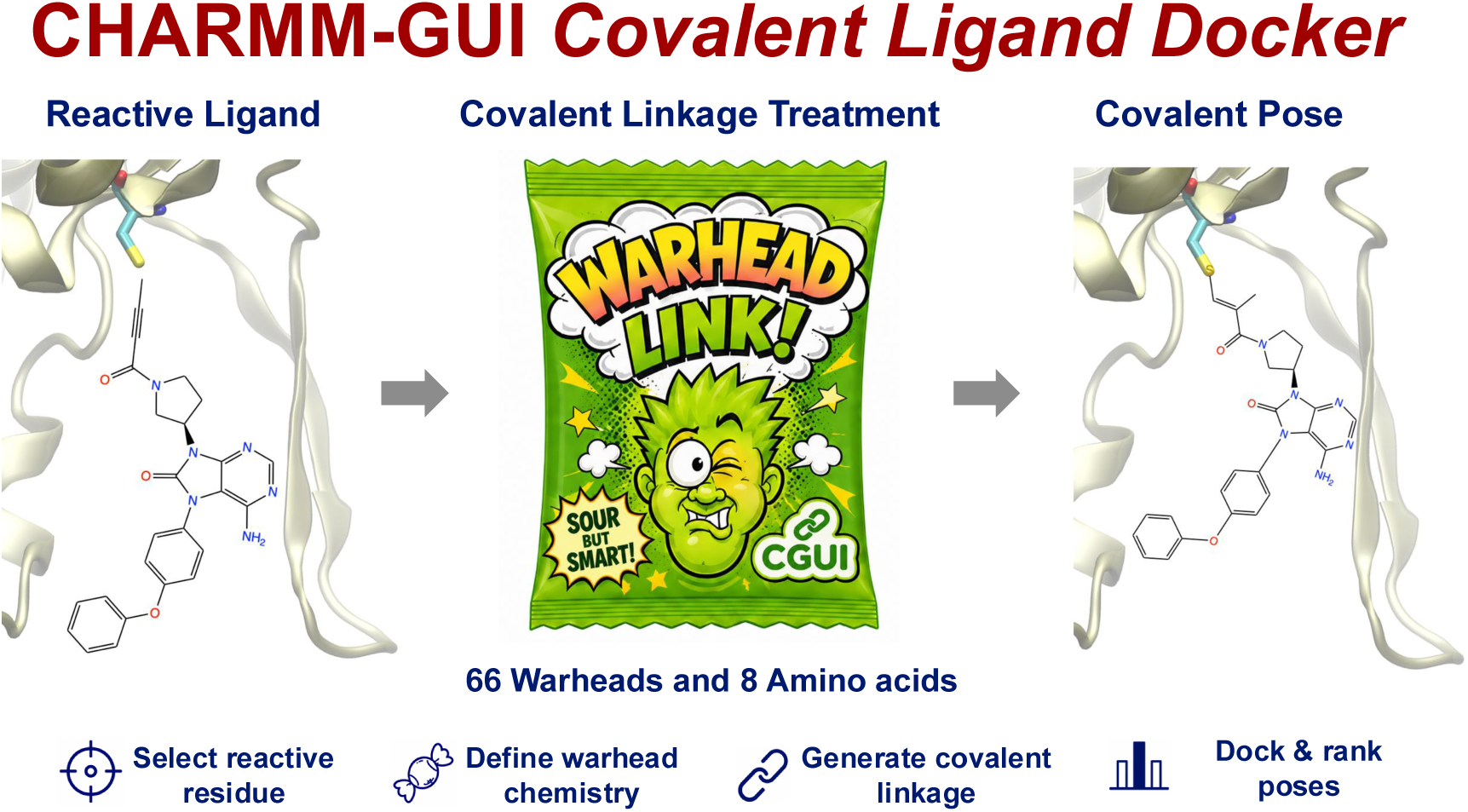

## Introduction

Covalent inhibitors, which form covalent bonds to target proteins, have gained increasing attention with advantages in prolonged binding, leveled potency, and reduced drug resistance.^1–6^ Historically, concerns regarding idiosyncratic adverse effects limited enthusiasm for developing covalent inhibitors. Nevertheless, many clinically important drugs, including penicillin and aspirin, were later found to act through covalent mechanisms after their discovery by serendipity.^1,7,8^ More recently, advances in screening technologies and rational design strategies, including structure-based drug design, have enabled researchers to fine-tune covalent mechanisms to mitigate potential liabilities and optimize pharmacological profiles.^1,4,5,8,9^

The mode of action of a covalent ligand is commonly described by a two-step model. In the first step, the ligand forms a reversible non-covalent complex with the target protein. In the second step, covalent bond formation occurs, transforming the pre-reaction ligand into a post-reaction adduct. The concept of targeted covalent inhibition combines a non-covalent binding scaffold with a reactive electrophilic warhead that covalently binds to a poorly conserved residue in the target protein. More than 70 new warheads have been discovered in the recent decades, and the exploration and design of new warheads continue to evolve rapidly.^3,7,10^ Consistent with this growth, publications and citations related to covalent inhibitors have increased sharply, with thousands of publications appearing each year.^2^ In addition, more than 25,000 protein–covalent ligand complexes are available in the RCSB PDB^11^ as of June 2026.

Computational approaches for covalent ligand research have also expanded rapidly. Commonly used approaches include molecular docking, molecular dynamics (MD), quantum mechanics (QM), machine learning (ML), and artificial intelligence (AI), which are used to predict the structures, energetics, and interactions of covalent ligands with target proteins.^9,12–15^ Databases such as CovPDB,^16^ CovBinderInPDB,^17^ and CovalentInDB 2.0^18,19^ serve as well-organized platforms containing curated information on covalent inhibitors and are valuable resources for structure-based virtual screening.

Molecular docking of covalent ligands, commonly referred to as covalent docking, is a widely used approach for rapidly evaluating covalent ligand binding poses. Current docking tools, including GOLD,^20–22^ CovDock,^23^ ICM-Pro,^24^ GNINA,^25^ and AutoDock4^26^ support covalent docking. However, despite the availability of multiple covalent docking tools, significant practical challenges remain. These include restrictions associated with commercial licenses, the level of expertise required to use specialized software, and the tedious preparation of covalent ligand structures. In particular, preparing a covalent ligand in its post-reaction adduct form, which is often required as input for covalent docking, can be time-consuming and nontrivial.

CHARMM-GUI (https://charmm-gui.org),^27^ a web-based cyberinfrastructure whose development was initiated in 2006, is dedicated to facilitating complex molecular system building by reducing expertise barriers and workload, allowing users to focus on the scientific questions and goals. CHARMM-GUI is actively maintained and continuously expanded with new functional modules. In a previous update, CHARMM-GUI introduced support for covalent ligand modeling and MD simulations through *PDB Reader & Manipulator*, the core module for reading and modifying biomolecular structures.^15,28^

In this work, we present *Covalent Ligand Docker* (CGUI-CLD), a new module that enables users to perform covalent docking through a semi-automated workflow using AutoDock4,^24^ together with extensive benchmark testing. A key feature of CGUI-CLD is its built-in knowledge-based library, which enables seamless transformation of covalent ligands into docking-ready adduct forms. Moreover, CGUI-CLD is connected to CHARMM-GUI *High-Throughput Simulator* (CGUI-HTS)^29,30^ to generate MD simulation systems and inputs for docked complexes, allowing users to further validate and screen docking results. Video demos for CGUI-CLD are available at https://www.charmm-gui.org/demo/covdock/1 to assist users with proper setup and usage.

## Methods

### Workflow of Covalent Ligand Docker

CGUI-CLD supports covalent docking with AutoDock4,^26,31^ which was selected for its robustness, performance, and license accessibility. AutoDock4 provides two covalent docking algorithms: the two-point attractor method and the flexible side-chain method, among which the latter shows significantly better performance.^26^ Therefore, we only support the flexible side-chain method in CGUI-CLD. In the flexible side-chain method, the target residue and ligand are combined into a single structure file. The protein backbone atoms are kept rigid, whereas the target side-chain atoms and ligand atoms are treated as flexible with conformational sampling performed through torsional rotations.

CGUI-CLD is seamlessly integrated with *PDB Reader & Manipulator*, which allows users to read and manipulate a biomolecular structure with options such as protonation, mutation, missing residue modeling, and disulfide-bond assignment.^15^ After a PDB structure is uploaded and, if necessary, modified, users can either select a bound ligand from the PDB file or upload ligand structures in SDF format to prepare ligand-residue structures. On the setup page (**Figure S1**), cofactor selection and ligand preparation are performed with 3D visualization using NGL viewer.^32^ Considering the diverse warheads and reaction mechanisms, preparing such ligand-residue structures in adduct form is a nontrivial work. To facilitate this process, CGUI-CLD provides an intuitive pop-up interface for each ligand, supported by the Marvin JS sketchpad,^33^ to guide ligand preparation (**Figure 1**). This interface requires two steps. In Step 1, users specify the target residue using a drop-down menu. In Step 2, users construct the ligand-residue structure, and CGUI-CLD supports both pre-reaction ligand and post-reaction ligand cases.

**Figure 1.**
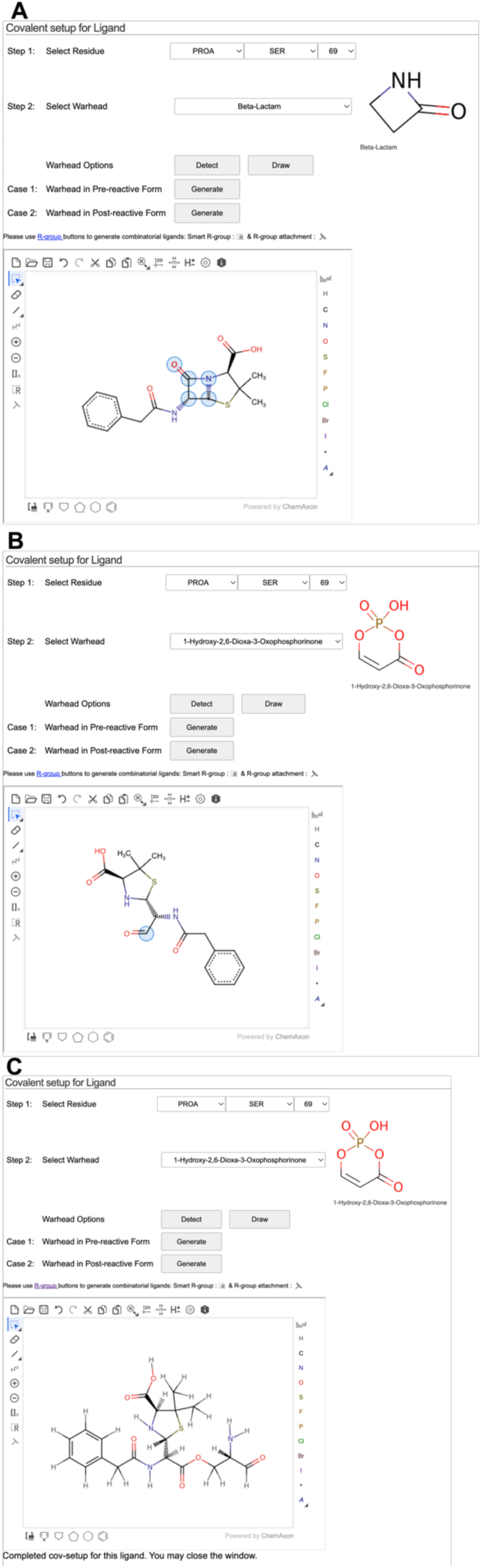
User interface for covalent ligand preparation of penicillin G. The first step in this example is to select Ser69 as the target residue in Step 1. (A) For preparation from the pre-reaction form of penicillin, users need to select the corresponding warhead, highlight the warhead atoms in the Marvin JS sketchpad manually or by using “Detect” button, and click “Case 1: Generate.” (B) For preparation from the post-reaction form of penicillin, users highlight the linkage atom and click “Case 2: Generate.” (C) The same penicillin-serine linkage structure is generated from both Case 1 and Case 2.

For the pre-reaction ligand case (**Figure 1A**), users need to upload either a pre-reaction ligand or a non-covalent ligand scaffold. If a non-covalent ligand scaffold is uploaded, CGUI-CLD provides an option to add a selected warhead from the drop-down menu directly onto the sketchpad using the “Draw” button. If a pre-reaction ligand containing a warhead is uploaded, the warhead must be selected to continue. In some cases, warheads such as epoxides or aziridines, contain symmetric ring structures (e.g., **Figure 3C**), so that users need to manually delete a bond to specify the bond that breaks during the ring-opening reaction. After modifying the ligand, users highlight the warhead atoms in the sketchpad according to the reference image shown in the upper-right corner of the webpage. This can be done either manually or by clicking the “Detect” button, which provides a convenient automated option. Users then click “Case 1: Generate,” after which CGUI-CLD prepares the ligand-residue structure and displays it in the sketchpad (**Figure 1C**). CGUI-CLD uses a built-in library, supported by RDKit,^34^ to detect and transform the warhead according to the warhead type, target residue, and reaction chemistry: a total of 104 warhead-residue combinations with 66 warheads and 8 amino acid residues (**Table S1**). Leaving groups, when applicable, are also removed automatically during this workflow.

For the post-reaction ligand case (**Figure 1B**), Step 2 requires users to highlight only one ligand atom that should be connected to the target residue side chain. Users then click “Case 2: Generate,” and the prepared ligand-residue structure is displayed in the sketchpad (**Figure 1C**). This option for ligand preparation can be used even when the corresponding warhead is not supported in the CGUI-CLD built-in library.

After ligand-residue structures are prepared, CGUI-CLD post-processes them through a unified workflow that follows the standard AutoDock4 procedure.^26^ RDKit^34^ is used to detect and rename backbone atoms and to align the prepared structure to the original target residue backbone coordinates. PDBQT files, grid parameters, and docking parameters are then generated using AutoDockTools.^31^ After covalent docking using AutoDock4, CGUI-CLD displays the docking score and pose rankings with real-time visualization for each pose (**Figure S2**). Users can download related files, including modified PDB file, pose coordinates, and ligand-residue input structures for further inspection. Selected poses can be transferred to CGUI-HTS for high-throughput MD simulation system setup.

### Computational details for benchmark test

The benchmark dataset reported by Scarpino et al.^35^ was used to evaluate CGUI-CLD. Thus, 207 protein-ligand complexes were covalently docked with AutoDock4 using CGUI-CLD. While the CGUI-CLD default option is to generate 10 poses for each protein-ligand case, the actual number of poses could be less than 10, depending on the covalent ligand conformation space in each binding site. For 207 protein-ligand complexes, CGUI-CLD AutoDock4 produced 865 poses in total. These poses were then transferred to CGUI-HTS for automated generation of MD simulation system and input files.

The CHARMM36m^36^ force field was used for the proteins, and CGenFF^37^ was used for the ligands, following the covalent ligand parameterization workflow implemented in *PDB Reader & Manipulator*.^15^ Each protein-ligand complex was solvated in 0.15 M KCl. Long range electrostatic interactions were calculated using the particle-mesh Ewald method,^38^ and van der Waals interactions were treated with a force-switching function between 10 Å and 12 Å.^39^ Hydrogen atoms were constrained with the SHAKE algorithm.^40^ Hydrogen mass repartitioning^41^ was applied to enable a 4-fs time step. Simulations were performed in the NPT (constant particle number, pressure, and temperature) ensemble at 303.15 K, and 10-ns production simulations were conducted using OpenMM^42^ with input files generated by CHARMM-GUI.^25^ The final snapshot from each MD trajectory, referred to as the MD-refined pose, was extracted for subsequent ligand RMSD and MM/GBSA analysis.

### Ligand RMSD and MM/GBSA

For both docking poses and MD-refined poses, protein structures were first aligned to their corresponding native structures. Ligand RMSD values relative to the native ligand conformations were then calculated to evaluate pose accuracy. In addition, ligand RMSD values between MD-refined poses and their corresponding initial docking poses were also calculated to assess pose stability during MD simulations.

MM/GBSA calculations were performed using OpenMM with the OBC2 GB model.^43^ The solvent and solute dielectric constants were set to 78.5 and 1, respectively, and the salt concentration was set to 0.15 M. The ligand binding free energy (ΔG_bind_) was estimated as:

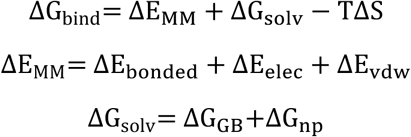

where ΔE_MM_and ΔG_solv_ are the changes in molecular mechanics energy and solvation free energy upon ligand binding, and the entropic contribution (TΔS) was neglected in this study. Thus, the overall energy terms include electrostatic (ΔE_elec_), van der Waals (ΔE_vdw_), GB electrostatic solvation (ΔG_GB_), and nonpolar solvation (ΔG_np_). The solvent-accessible surface area (SASA) was measured with a probe radius of 1.4 Å to estimate the nonpolar solvation energy with a surface tension coefficient (*γ*) of 0.0072 kcal/mol/Å^2^: ΔG_np_ = *γ* · SASA.

### Analysis of pose classifiers

Receiver operating characteristic (ROC) analysis was performed to evaluate potential classifiers for distinguishing correct and incorrect binding poses. For fair comparisons among poses generated for the same protein-ligand case, relative docking scores were calculated as the difference between the docking score of each pose and that of the Top-1 scored docking pose for the corresponding case. Similarly, relative MM/GBSA energies were calculated as the difference between the MM/GBSA energy of each pose and that of the Top-1 MM/GBSA-ranked pose for the corresponding case. The optimal cutoff for each classifier was determined by maximizing Youden’s index, defined as the difference between the true positive rate and the false positive rate.

## Results and Discussion

### Warhead coverage and built-in library statistics in CGUI-CLD

The CGUI-CLD built-in library has been established based on the protein-ligand complexes from CovPDB^16^ and CovBinderInPDB^17^ by selecting each unique warhead-residue combination and the algorithm was tested if a ligand can be correctly transformed to the ligand-residue linkage structure. CGUI-CLD currently supports a total of 104 warhead-residue combinations, covering 66 warheads and 8 amino acid residues (**Table S1**). As expected, cysteine and serine residues are the two most popular target residues covered by CGUI-CLD, supported by 45 warheads and 25 warheads, respectively (**Figure 2**). The top popular warheads in CovPDB^16^, such as vinyl carbonyl, beta-lactam, ketone, boronic acid, and halomethyl carbonyl, are also covered. Some rare warheads are not supported due to unclassified reaction mechanisms. In case of unsupported warheads, we recommend the post-reaction ligand option (**Figure 1B**) in which users can upload or draw the ligand adduct directly. Overall, CGUI-CLD supports most popular cases in covalent docking and provides flexibility for ligand structure preparation.

**Figure 2.**
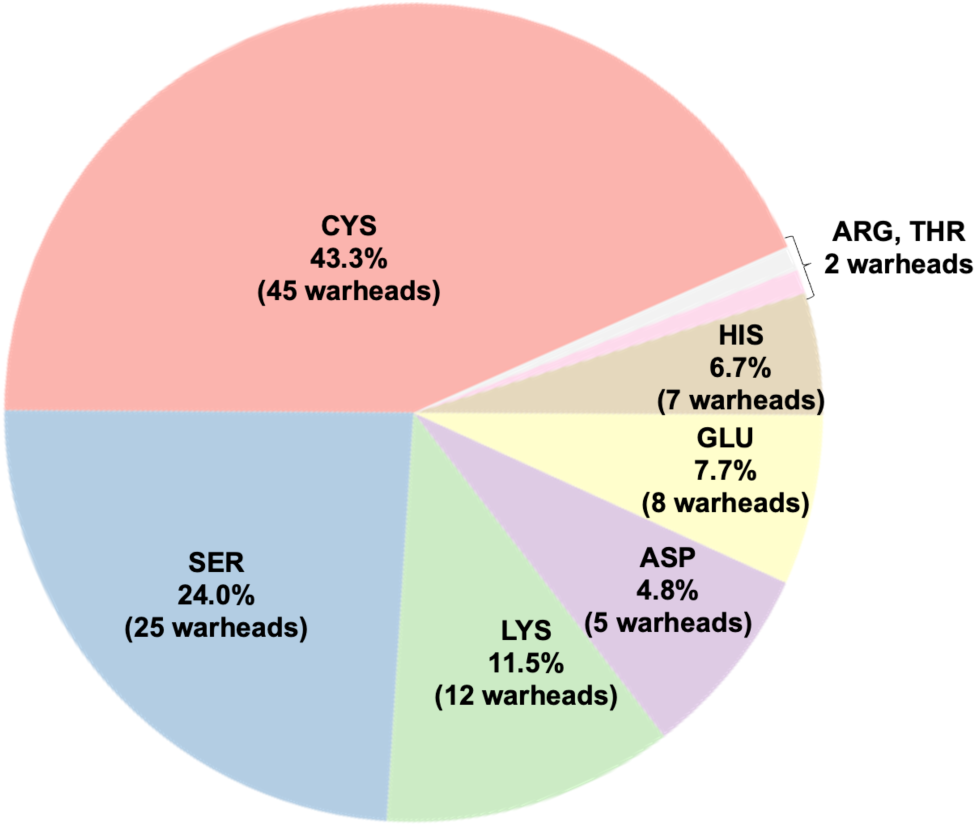
Distribution of amino acids in the warhead-amino acid combinations available in the CGUI-CLD built-in library for automated detection: a total of 104 warhead-residue combinations with 66 warheads and 8 amino acid residues.

### Covalent docking case studies

We present three specific covalent docking cases to illustrate CGUI-CLD usages in the tutorials. The top-scored pose was selected to investigate molecular interactions between the protein and the ligand. The covalent reaction for each case is depicted in **Figure 3**.

**Figure 3.**
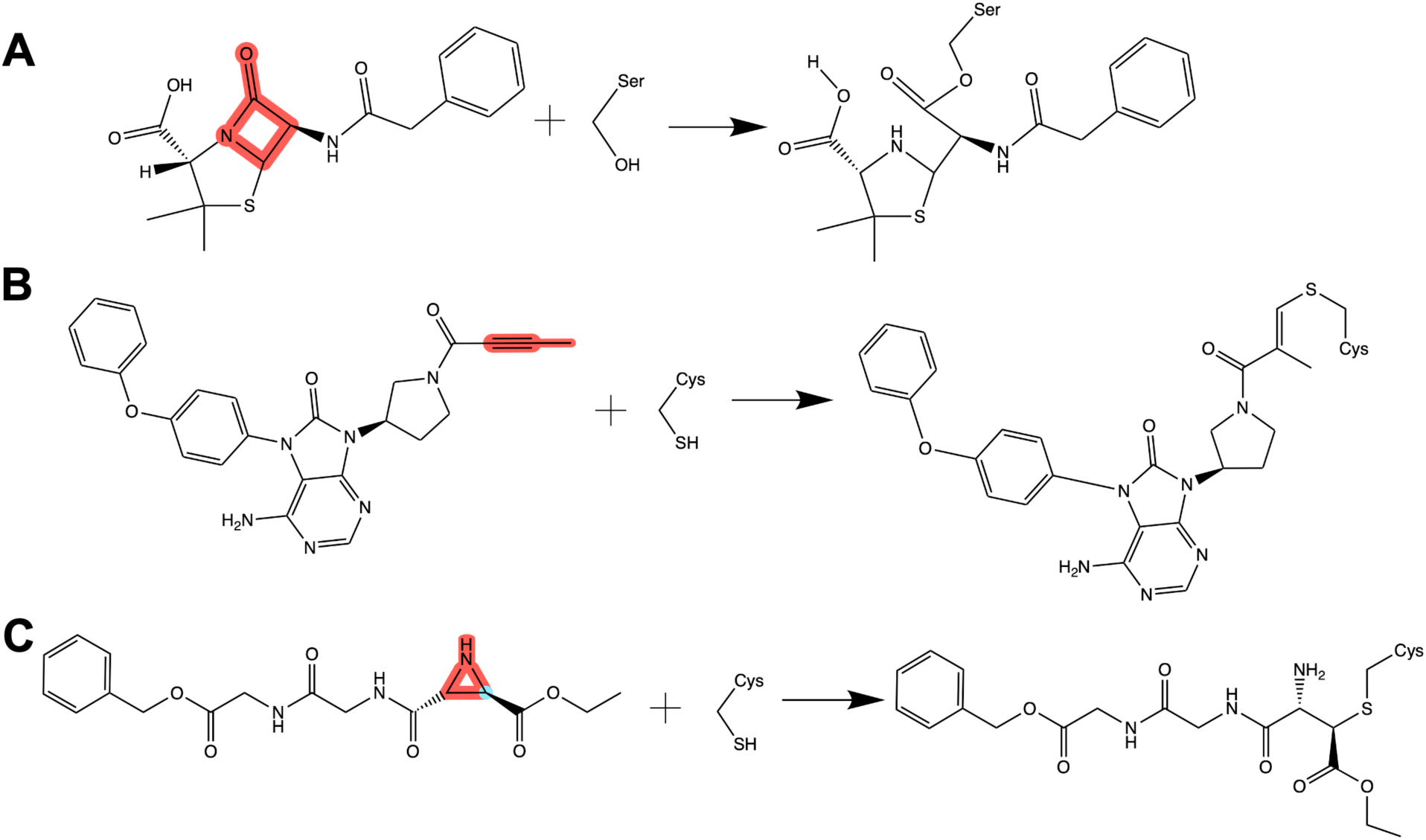
Illustrations of covalent reactions in example docking cases. The covalent warheads are highlighted in red. The adduct forms of ligand-residue structure shown on the right side is required for covalent docking. (A) Penicillin G reacts with serine in PDB ID 5KMW. (B) Tirabrutinib reacts with cysteine in PDB ID 5P9J. (C) A trans-aziridine ligand reacts with cysteine in PDB ID 2A5I. The electrophilic carbon (in cyan) is subject to a nucleophilic attack.

Penicillin is the most popular antibiotic drug, discovered by serendipity in 1928 and changed medical history.^44^ It features a ring-opening reaction and is covalently linked to the catalytic serine of penicillin-binding proteins (**Figure 3A**). The crystal structure of Toho1 beta-lactamase in complex with penicillin (PDB ID 5KMW) was published in 2016, with penicillin G bound to Ser69 in the catalytic site.^45^ In this case, our covalent docking of penicillin onto Ser69 shows an RMSD of 0.71 Å from the native ligand structure (**Figure 4A**). Notably, the sidechains of Ser236 and Thr234 form hydrogen bonds with the ligand, and the amide groups of Asn103 and Asn131 also interact with the ligand, respectively. These interactions are considered crucial in stabilizing penicillin during the hydrolysis step, and the Top-1 docking pose is consistent with the experimental results.

**Figure 4.**
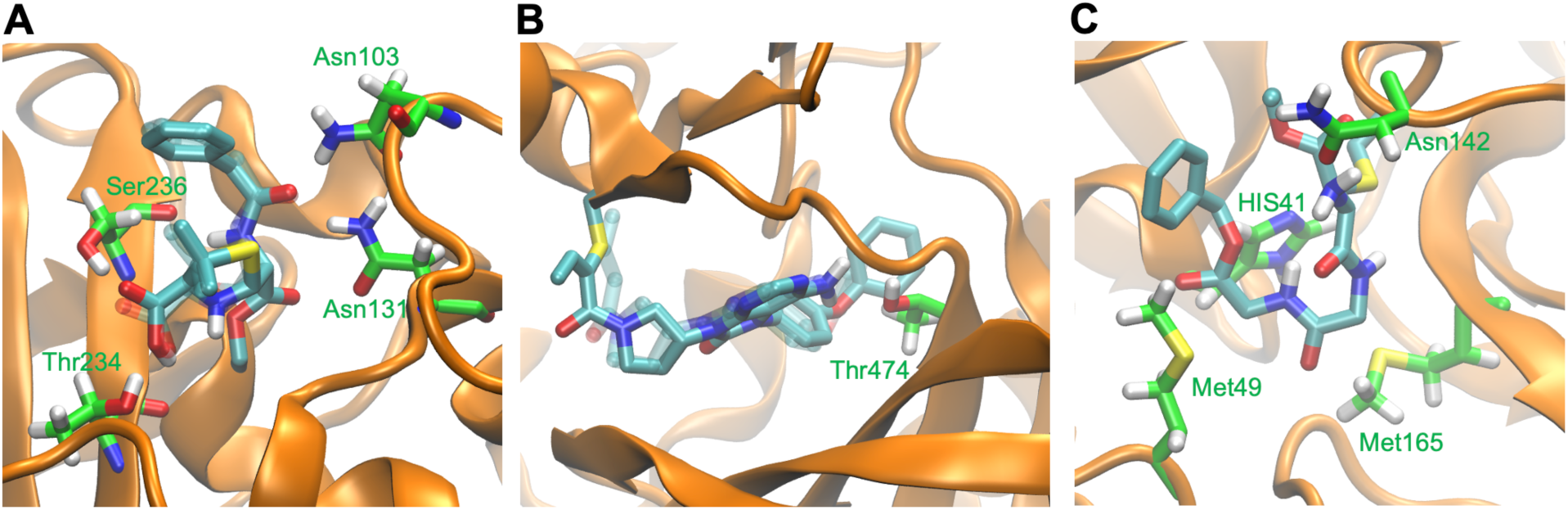
Snapshots of Top-1 docking pose in three test cases: PDB ID (A) 5KMW, (B) 5P9J, and (C) 2A5I. Ligands are shown as sticks with cyan carbon, and interacting residues are shown as sticks with green carbon. The reference native pose (in (A) and (B)) is shown in transparent sticks.

Bruton Tyrosine Kinase (BTK) is an oncoprotein and has been a popular target to treat B-cell malignancies or autoimmune diseases.^46^ Ibrutinib, an FDA-approved irreversible inhibitor, was resolved in complex with the human BTK kinase domain (PDB ID 5P9J) and found to be covalently attached to Cys481 at the catalytic site.^47^ Tirabrutinib, as an analogue of Ibrutinib, features a butynamide warhead and a 6-5 membered heterocyclic ring as shown in **Figure 3B** (PDB ID 5P9M: Tirabrutinib in complex with human BTK). To test covalent cross-docking, Tirabrutinib was docked onto Cys481 in PDB 5P9J. After superimposing the structure onto PDB 5P9M,^47^ the Top-1 docking pose yields an RMSD of 1.70 Å with respect to the native ligand structure, with the phenoxy-phenyl group towards the hydrophobic pocket (**Figure 4B**). Hydrogen bonds are found between the ligand and Thr474, but missed with Met477 and Glu475, which may be due to the limitation of rigid docking on a different protein conformation (i.e., induced fit effects). Our results suggest the limitation of cross-docking that is still an underexplored aspect in covalent docking.

SARS-CoV main peptidase (M^pro^) is the major target for treating severe acute respiratory syndrome. Covalent inhibitors were designed to target Cys145 in the catalytic site of SARS-CoV M^pro^, with a warhead such as α-ketoamide, epoxide, or α-fluoroketone.^48,49^ Previously, an aza-peptide epoxide inhibitor in complex with SARS-CoV M^pro^ (PDB ID 2A5I) was resolved, and the inhibitor is well-positioned inside the substrate binding pocket.^50^ Also, a trans-aziridine inhibitor (**Figure 3C**) was confirmed to covalently inhibit SARS-CoV M^pro^ through HPLC-based assay, with 54% inhibition at 100 μM.^51^ We docked the trans-aziridine inhibitor to Cys145 in PDB 2A5I. As shown in **Figure 4C**, the benzene moiety extends towards the S2 hydrophobic pocket and interacts with Met49, while the ethyl group fits inner side of the binding pocket with hydrophobic interactions. Asn142, HIS41, and Met165 also show interactions with the ligand. Despite no experimental covalent structure was resolved for this trans-aziridine inhibitor, a traditional non-covalent docking study was performed using FlexX,^51,52^ which showed a similar orientation of the ethyl group as in **Figure 4C**. There are also differences that could be contributed by the covalent complex formation or different docking parameters. For example, the pose in our case fits more into the S2 and S4 pockets, which is similar to a peptide-like chloromethyl ketone inhibitor structure (PDB ID 1UK4).^53^ Overall, our results show a reasonable conformation of the ligand in fitting the binding pocket of SARS-CoV M^pro^.

### Benchmark testing on covalent docking and high-throughput MD simulations

To systematically evaluate AutoDock4 performance using CGUI-CLD and explore the potential of MD refinement on binding poses using CGUI-HTS, we used the 207 protein-ligand complex dataset from Scarpino et al.^35^ and conducted a benchmark study. For each complex case, AutoDock4 produced different numbers of poses (with a maximum number of 10) and **Figure S3** shows a distribution of how many cases produced how many poses. For example, 31 protein-ligand complex cases produced only 1 pose cluster and only 1 protein-ligand case yielded 10 poses, indicating much less ligand conformational space for covalent docking compared to typical noncovalent docking. In total, there are 865 poses from the 207 protein-ligand complex cases. Our docking results were categorized by best AutoDock4-scored poses (Top-1 pose), best RMSD poses, and all available poses.

Among Top-1 poses, 18.4% are within 1 Å RMSD from the native conformations, 27.1% within 1-2 Å, and 34.8% within 2-3 Å (**Figure 5A**). Thus, the prediction success rate is 18.4%, 45.5%, and 80.3% when the RMSD cutoff is 1 Å, 2 Å, and 3 Å, respectively. Compared with the result from Scarpino et al. (16%, 55%, 68%), the deviations are expected due to the random seeds in sampling. With best docking RMSD poses, the cumulative success rates are 22.7%, 69.6%, and 94.7% with an RMSD cutoff of 1 Å, 2 Å, and 3 Å, respectively, which is close to the best docking RMSD pose result from Scarpino et al. (30%, 75%, and 87%). Overall, these results show the consistency of AutoDock4 performance using CGUI-CLD.

**Figure 5.**
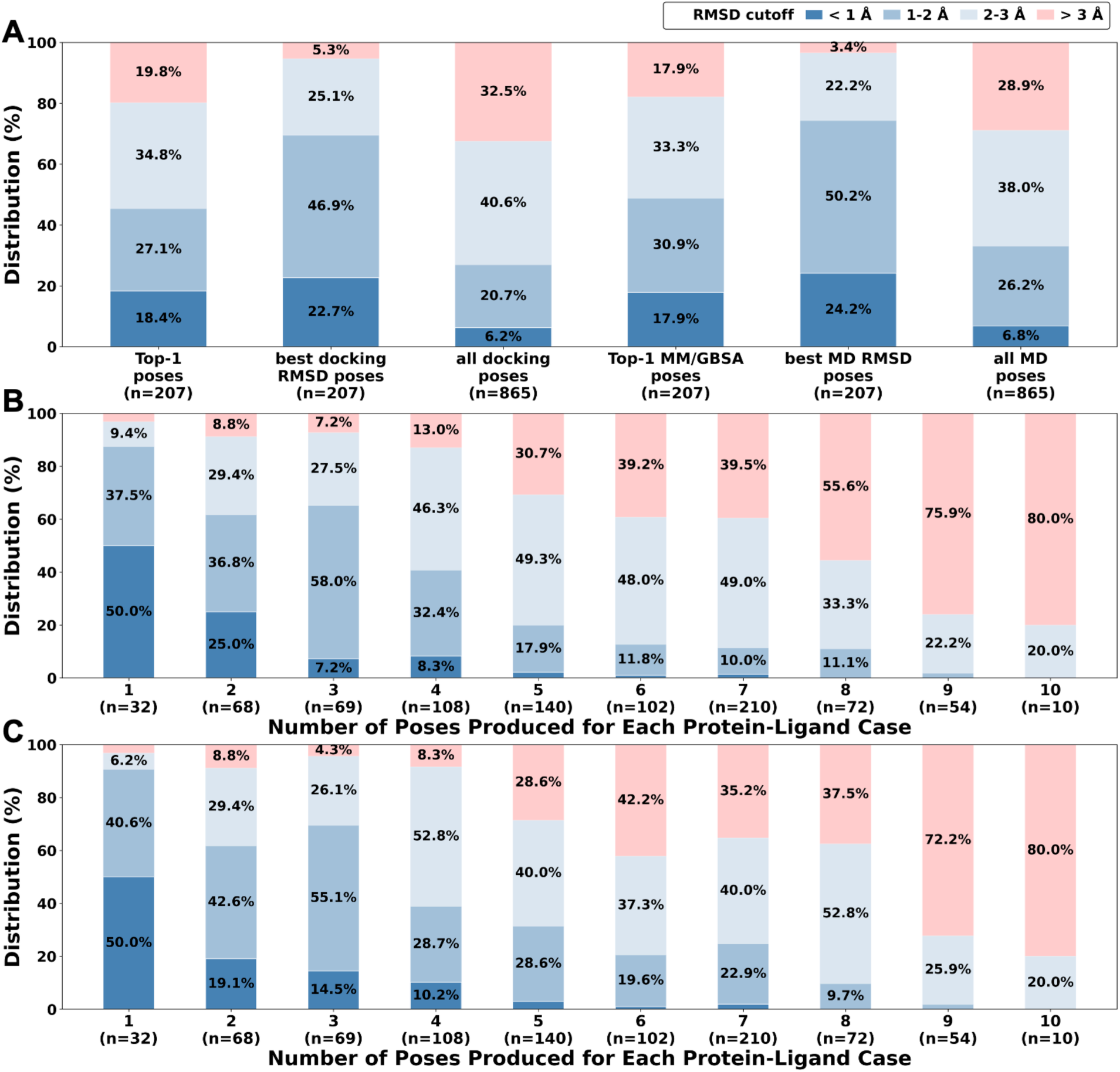
Distribution of ligand RMSD to the native structures using the dataset from Scarpino et al. (A) Overall distributions of Top-1 poses, best docking RMSD poses, all docking poses, Top-1 MM/GBSA poses, best MD RMSD poses, all MD poses. (B) All ligand RMSD distributions grouped with the same number of poses in docking. (C) All ligand RMSD distributions grouped with the same number of poses after MD simulations.

The sampling power of AutoDock4 is further explored by analyzing the success rate across all poses. Among all 865 poses, only 6.2% are successfully retrieved in near-native conformations within 1 Å, as shown in **Figure 5A**. This number substantially increases to 67.9% with an RMSD cutoff of 3 Å. The RMSD distributions from all poses are then split by the same number of poses produced for each docking case (**Figure 5B**). The protein-ligand complex cases that produced only 1 pose (i.e., 31 cases in **Figure S3**) show the highest success rate (50.0% and 96.9% within 1 Å and 3 Å, respectively), while for the cases with 5 poses, the success rate drops to 2.1% and 69.3%. For the cases with 9 poses, the success rate further drops to 24.1% even with a 3 Å cutoff, meaning only 13/54 poses were successfully sampled. This trend is expected as covalent ligands with higher degrees of freedom (with more poses) are more difficult to be scored correctly. It is commonly recognized that a docking score may not be sufficient in evaluating binding poses. In our results, incorrect binding poses (native RMSD > 2.5 Å) are also largely distributed with high scores (**Figure S4**).

To explore if short MD simulations can improve binding poses by examining the pose stability, we used all 865 poses as the starting structures and performed 10-ns MD simulations using CGUI-HTS. In addition, we performed the MM/GBSA analysis as a metric to re-rank poses after MD simulations. 10-ns MD trivially increases success rate across Top-1 MM/GBSA poses, best MD RMSD poses, and all MD poses (**Figure 5A**). Throughout the simulation, compared with correct poses (native RMSD <= 2.5 Å), incorrect binding poses show a trend of instability with higher ligand RMSD and a more even distribution in relative MM/GBSA differences to the Top-1 MM/GBSA poses (**Figure 6A,B**).

**Figure 6.**
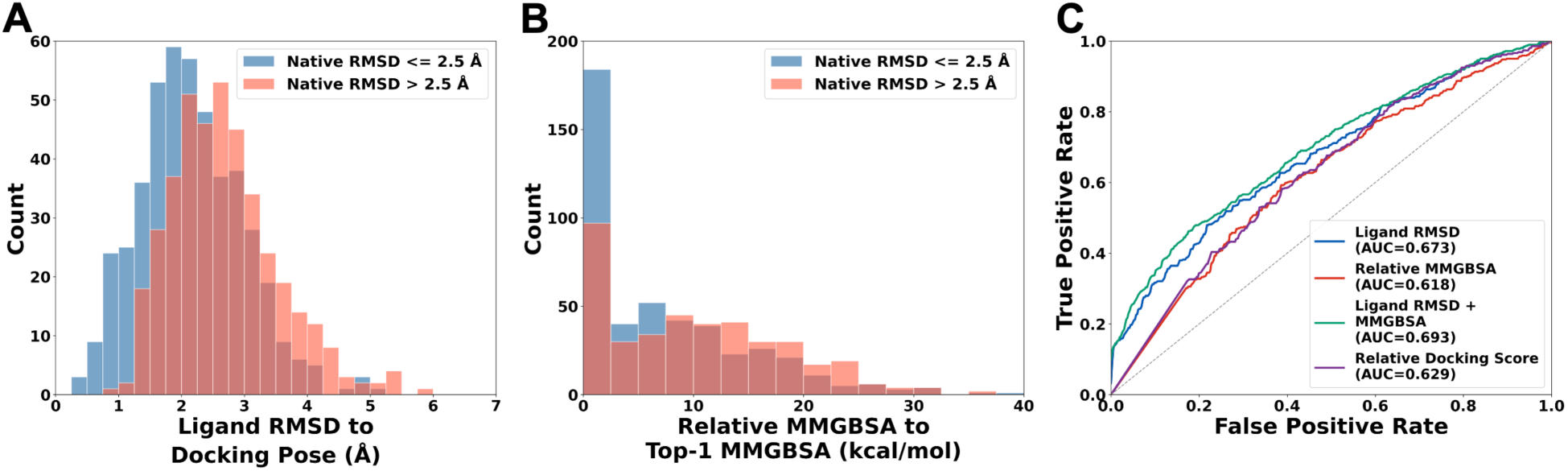
Histogram distributions of (A) ligand RMSD to their docking poses after MD simulations and (B) relative MM/GBSA difference to the Top-1 MM/GBSA in each testcase. The histograms include 865 poses. The ligand poses are categorized based on its native RMSD): blue for native RMSD <= 2.5 Å (n = 461) and red for native RMSD > 2.5 Å (n = 404). (C) The ROC plot shows the performance of various classifiers: ligand MD RMSD to their docking poses (in blue), relative MM/GBSA (in red), ligand MD RMSD + relative MM/GBSA (in green), and relative docking scores (in purple).

We then evaluated the performance of several metrics (relative docking scores, ligand MD RMSD to initial docking poses, relative MM/GBSA) in distinguishing incorrect poses from correct ones (**Figure 6C**). The ROC analysis shows moderate performance in relative docking score (AUC=0.629) and relative MM/GBSA (AUC=0.618), while ligand MD RMSD (AUC=0.673) and ligand MD RMSD + relative MM/GBSA (AUC=0.693) exhibit improved performance compared to relative docking scores (9% and 13% improvement, respectively). Based on Youden-index, the optimal cutoff is 2.04 Å for ligand MD RMSD, 2.15 Å + 13.6 kcal/mol for ligand MD RMSD + relative MM/GBSA, while 1.3 kcal/mol and 7.3 kcal/mol for relative docking scores and relative MM/GBSA, respectively. Previously, it was reported that short MD simulations can moderately improve the binding poses for traditional noncovalent ligands.^30^ Our work shows that short MD simulations are also applicable in refining binding poses for covalent ligands, and suggests that the MD metrics (ligand RMSD and MM/GBSA) can be helpful as pose classifiers in covalent ligand screening.

### Conclusions

Despite the rapid evolution of covalent warheads discovery and increasing number of resolved covalent protein-ligand complex structures each year, covalent docking is still a growing field. Previously, Scarpino et al. performed a comprehensive evaluation of covalent docking tools, which shows that AutoDock4 is balanced in performance and computational cost, among popular covalent docking tools.^35^ More docking parameters and methodologies are currently tested and applied on covalent ligands. For example, CovCIFDock utilizes QM/MM potentials to include reaction energetics in their scoring function.^54^ HCovDock also includes a covalent bond-based energy correction in docking simulation.^55^ Beyond traditional covalent docking tools, Shamir et al.^56^ recently discovered a superior performance of AlphaFold3 in predicting covalent ligand poses in a co-folding manner.

Preparation of ligands in covalent docking, especially transforming covalent ligands to their adduct state, is a tedious and error-prone process. A few popular docking programs, such as Schrodinger CovDock,^23^ provides a default covalent reaction repository, yet the number of warheads is still limited to a dozen. A custom reaction file is required if a desired warhead type is not available. Cavities 2.0^57^ shows an excellent docking performance and supports 15 types of warheads, but also requires command line implementation and long computational time.

In this work, we present CHARMM-GUI *Covalent Ligand Docker* as a web-based covalent docking platform for AutoDock4 with a user-friendly interactive interface and seamlessly integrate it with CGUI-HTS for all-atom MD simulation setup. Currently, CGUI-CLD provides the most diverse selection of warheads (66 warheads and 8 amino acids), support smooth transformation of covalent ligands to their adduct state, and features a visualized sketchpad for easy inspection and manipulation of ligands. This is motivated by ongoing discovery of increasing number of warheads and the demand for a user-friendly interface. To the best of our knowledge, this is the first free online covalent docking platform that supports the full preparation of ligands. Also, our workflow, especially the ligand transformation function, could be generalized, since the ligand adduct structure is the direct input for most covalent docking programs. We expect to expand CGUI-CLD by supporting more covalent docking algorithms and warheads in the future. It is our hope that CGUI-CLD can boost the progress of structure-based virtual screening on covalent ligands and contribute to the covalent ligand research.

## Supporting Information

Additional details with CGUI-CLD built-in library, representative screenshots, and benchmark docking analyses, including pose clusters and docking scores.

## Author Contributions

L.K.: software development, docking and MD simulations, analysis, visualization, and writing— original draft, review and editing; D.S.: software development, visualization, and writing—original draft, review and editing; and W.I.: conceptualization, supervision, and writing—review and editing.

## Acknowledgments

This work is supported by NIH R35 GM153458.

## DATA AND SOFTWARE AVAILABILITY

The data underlying this study, including the input files for model system generation are freely available in *Ligand Docker* Archive in CHARMM-GUI (http://www.charmm-gui.org/docs/archive/covligdock), and can be reproduced in CHARMM-GUI *Ligand Docker* (https://charmm-gui.org/input/ligdock).

## References

(1) Singh, J.; Petter, R. C.; Baillie, T. A.; Whitty, A. The Resurgence of Covalent Drugs. Nat. Rev. Drug Discov. 2011, 10 (4), 307–317. 10.1038/nrd3410.

(2) Sutanto, F.; Konstantinidou, M.; Dömling, A. Covalent Inhibitors: A Rational Approach to Drug Discovery. RSC Med. Chem. 2020, 11 (8), 876–884. 10.1039/D0MD00154F.

(3) Gehringer, M.; Laufer, S. A. Emerging and Re-Emerging Warheads for Targeted Covalent Inhibitors: Applications in Medicinal Chemistry and Chemical Biology. J. Med. Chem. 2019, 62 (12), 5673–5724. 10.1021/acs.jmedchem.8b01153.

(4) Schaefer, D.; Cheng, X. Recent Advances in Covalent Drug Discovery. Pharmaceuticals 2023, 16 (5), 663. 10.3390/ph16050663.

(5) Bauer, R. A. Covalent Inhibitors in Drug Discovery: From Accidental Discoveries to Avoided Liabilities and Designed Therapies. Drug Discov. Today 2015, 20 (9), 1061–1073. 10.1016/j.drudis.2015.05.005.

(6) Adeniyi, A. A.; Muthusamy, R.; Soliman, M. E. New Drug Design with Covalent Modifiers. Expert Opin. Drug Discov. 2016, 11 (1), 79–90. 10.1517/17460441.2016.1115478.

(7) Hillebrand, L.; Liang, X. J.; Serafim, R. A. M.; Gehringer, M. Emerging and Re-Emerging Warheads for Targeted Covalent Inhibitors: An Update. J. Med. Chem. 2024, 67 (10), 7668–7758. 10.1021/acs.jmedchem.3c01825.

(8) Boike, L.; Henning, N. J.; Nomura, D. K. Advances in Covalent Drug Discovery. Nat. Rev. Drug Discov. 2022, 21 (12), 881–898. 10.1038/s41573-022-00542-z.

(9) Awoonor-Williams, E.; Walsh, A. G.; Rowley, C. N. Modeling Covalent-Modifier Drugs. Biochimica et Biophysica Acta (BBA) - Proteins and Proteomics 2017, 1865 (11), 1664–1675. 10.1016/j.bbapap.2017.05.009.

(10) Mehta, N. V.; Degani, M. S. The Expanding Repertoire of Covalent Warheads for Drug Discovery. Drug Discov. Today 2023, 28 (12), 103799. 10.1016/j.drudis.2023.103799.

(11) Burley, S. K.; Bhatt, R.; Bhikadiya, C.; Bi, C.; Biester, A.; Biswas, P.; Bittrich, S.; Blaumann, S.; Brown, R.; Chao, H.; Chithari, V. R.; Craig, P. A.; Crichlow, G. V.; Duarte, J. M.; Dutta, S.; Feng, Z.; Flatt, J. W.; Ghosh, S.; Goodsell, D. S.; Green, R. K.; Guranovic, V.; Henry, J.; Hudson, B. P.; Joy, M.; Kaelber, J. T.; Khokhriakov, I.; Lai, J.-S.; Lawson, C. L.; Liang, Y.; Myers-Turnbull, D.; Peisach, E.; Persikova, I.; Piehl, D. W.; Pingale, A.; Rose, Y.; Sagendorf, J.; Sali, A.; Segura, J.; Sekharan, M.; Shao, C.; Smith, J.; Trumbull, M.; Vallat, B.; Voigt, M.; Webb, B.; Whetstone, S.; Wu-Wu, A.; Xing, T.; Young, J. Y.; Zalevsky, A.; Zardecki, C. Updated Resources for Exploring Experimentally-Determined PDB Structures and Computed Structure Models at the RCSB Protein Data Bank. Nucleic Acids Res. 2025, 53 (D1), D564–D574. 10.1093/nar/gkae1091.

(12) Singh, N.; Vayer, P.; Villoutreix, B. O. The Covalent Docking Software Landscape: Features and Applications in Drug Design. Brief. Bioinform. 2025, 26 (6). 10.1093/bib/bbaf697.

(13) Liu, R.; Vázquez-Montelongo, E. A.; Ma, S.; Shen, J. Quantum Descriptors for Predicting and Understanding the Structure–Activity Relationships of Michael Acceptor Warheads. J. Chem. Inf. Model. 2023, 63 (15), 4912–4923. 10.1021/acs.jcim.3c00720.

(14) Mihalovits, L. M.; Ferenczy, G. G.; Keserű, G. M. The Role of Quantum Chemistry in Covalent Inhibitor Design. Int. J. Quantum Chem. 2022, 122 (8). 10.1002/qua.26768.

(15) Kong, L.; Park, S.-J.; Im, W. CHARMM-GUI PDB Reader and Manipulator: Covalent Ligand Modeling and Simulation. J. Mol. Biol. 2024, 436 (17), 168554. 10.1016/j.jmb.2024.168554.

(16) Gao, M.; Moumbock, A. F. A.; Qaseem, A.; Xu, Q.; Günther, S. CovPDB: A High-Resolution Coverage of the Covalent Protein–Ligand Interactome. Nucleic Acids Res. 2022, 50 (D1), D445–D450. 10.1093/nar/gkab868.

(17) Guo, X.-K.; Zhang, Y. CovBinderInPDB: A Structure-Based Covalent Binder Database. J. Chem. Inf. Model. 2022, 62 (23), 6057–6068. 10.1021/acs.jcim.2c01216.

(18) Du, H.; Zhang, X.; Wu, Z.; Zhang, O.; Gu, S.; Wang, M.; Zhu, F.; Li, D.; Hou, T.; Pan, P. CovalentInDB 2.0: An Updated Comprehensive Database for Structure-Based and Ligand-Based Covalent Inhibitor Design and Screening. Nucleic Acids Res. 2025, 53 (D1), D1322– D1327. 10.1093/nar/gkae946.

(19) Du, H.; Gao, J.; Weng, G.; Ding, J.; Chai, X.; Pang, J.; Kang, Y.; Li, D.; Cao, D.; Hou, T. CovalentInDB: A Comprehensive Database Facilitating the Discovery of Covalent Inhibitors. Nucleic Acids Res. 2021, 49 (D1), D1122–D1129. 10.1093/nar/gkaa876.

(20) Verdonk, M. L.; Cole, J. C.; Hartshorn, M. J.; Murray, C. W.; Taylor, R. D. Improved Protein– Ligand Docking Using GOLD. Proteins: Structure, Function, and Bioinformatics 2003, 52 (4), 609–623. 10.1002/prot.10465.

(21) Cole, J.; Willem M. Nissink, J.; Taylor, R. Protein ÄìLigand Docking Virtual Screening with GOLD; 2005; pp 379–415. 10.1201/9781420028775.ch15.

(22) Jones, G.; Willett, P.; Glen, R. C.; Leach, A. R.; Taylor, R. Development and Validation of a Genetic Algorithm for Flexible Docking 1 1Edited by F. E. Cohen. J. Mol. Biol. 1997, 267 (3), 727–748. 10.1006/jmbi.1996.0897.

(23) Zhu, K.; Borrelli, K. W.; Greenwood, J. R.; Day, T.; Abel, R.; Farid, R. S.; Harder, E. Docking Covalent Inhibitors: A Parameter Free Approach To Pose Prediction and Scoring. J. Chem. Inf. Model. 2014, 54 (7), 1932–1940. 10.1021/ci500118s.

(24) Abagyan, R.; Totrov, M.; Kuznetsov, D. ICM—A New Method for Protein Modeling and Design: Applications to Docking and Structure Prediction from the Distorted Native Conformation. J. Comput. Chem. 1994, 15 (5), 488–506. 10.1002/jcc.540150503.

(25) Lee, J.; Cheng, X.; Swails, J. M.; Yeom, M. S.; Eastman, P. K.; Lemkul, J. A.; Wei, S.; Buckner, J.; Jeong, J. C.; Qi, Y.; Jo, S.; Pande, V. S.; Case, D. A.; Brooks, C. L.; MacKerell, A. D.; Klauda, J. B.; Im, W. CHARMM-GUI Input Generator for NAMD, GROMACS, AMBER, OpenMM, and CHARMM/OpenMM Simulations Using the CHARMM36 Additive Force Field. J. Chem. Theory Comput. 2016, 12 (1), 405–413. 10.1021/acs.jctc.5b00935.

(26) Bianco, G.; Forli, S.; Goodsell, D. S.; Olson, A. J. Covalent Docking Using Autodock: Two-point Attractor and Flexible Side Chain Methods. Protein Science 2016, 25 (1), 295–301. 10.1002/pro.2733.

(27) Jo, S.; Kim, T.; Iyer, V. G.; Im, W. CHARMM-GUI: A Web-based Graphical User Interface for CHARMM. J. Comput. Chem. 2008, 29 (11), 1859–1865. 10.1002/jcc.20945.

(28) Park, S.-J.; Kern, N.; Brown, T.; Lee, J.; Im, W. CHARMM-GUI PDB Manipulator: Various PDB Structural Modifications for Biomolecular Modeling and Simulation. J. Mol. Biol. 2023, 435 (14), 167995. 10.1016/j.jmb.2023.167995.

(29) Guterres, H.; Park, S.; Zhang, H.; Perone, T.; Kim, J.; Im, W. CHARMM-GUI *High-throughput Simulator* for Efficient Evaluation of Protein–Ligand Interactions with Different Force Fields. Protein Science 2022, 31 (9). 10.1002/pro.4413.

(30) Guterres, H.; Im, W. Improving Protein-Ligand Docking Results with High-Throughput Molecular Dynamics Simulations. J. Chem. Inf. Model. 2020, 60 (4), 2189–2198. 10.1021/acs.jcim.0c00057.

(31) Morris, G. M.; Huey, R.; Lindstrom, W.; Sanner, M. F.; Belew, R. K.; Goodsell, D. S.; Olson, A. J. AutoDock4 and AutoDockTools4: Automated Docking with Selective Receptor Flexibility. J. Comput. Chem. 2009, 30 (16), 2785–2791. 10.1002/jcc.21256.

(32) Rose, A. S.; Bradley, A. R.; Valasatava, Y.; Duarte, J. M.; Prlić, A.; Rose, P. W. NGL Viewer: Web-Based Molecular Graphics for Large Complexes. Bioinformatics 2018, 34 (21), 3755–3758. 10.1093/bioinformatics/bty419.

(33) Marvin JS Was Used for Drawing, Displaying and Characterizing Chemical Structures, Substructures and Reactions, Marvin JS 16.6.6, 2016, ChemAxon. Available at: Http://www.Chemaxon.Com.

(34) Landrum G. RDKit 2023.09.2 Documentation. https://www.rdkit.org/ 2023.

(35) Scarpino, A.; Ferenczy, G. G.; Keserű, G. M. Comparative Evaluation of Covalent Docking Tools. J. Chem. Inf. Model. 2018, 58 (7), 1441–1458. 10.1021/acs.jcim.8b00228.

(36) Huang, J.; Rauscher, S.; Nawrocki, G.; Ran, T.; Feig, M.; de Groot, B. L.; Grubmüller, H.; MacKerell, A. D. CHARMM36m: An Improved Force Field for Folded and Intrinsically Disordered Proteins. Nat. Methods 2017, 14 (1), 71–73. 10.1038/nmeth.4067.

(37) Vanommeslaeghe, K.; Hatcher, E.; Acharya, C.; Kundu, S.; Zhong, S.; Shim, J.; Darian, E.; Guvench, O.; Lopes, P.; Vorobyov, I.; Mackerell, A. D. CHARMM General Force Field: A Force Field for Drug-like Molecules Compatible with the CHARMM All-atom Additive Biological Force Fields. J. Comput. Chem. 2010, 31 (4), 671–690. 10.1002/jcc.21367.

(38) Darden, T.; York, D.; Pedersen, L. Particle Mesh Ewald: An *N* ⋅log(*N*) Method for Ewald Sums in Large Systems. J. Chem. Phys. 1993, 98 (12), 10089–10092. 10.1063/1.464397.

(39) Ryckaert, J.-P.; Ciccotti, G.; Berendsen, H. J. C. Numerical Integration of the Cartesian Equations of Motion of a System with Constraints: Molecular Dynamics of n-Alkanes. J. Comput. Phys. 1977, 23 (3), 327–341. 10.1016/0021-9991(77)90098-5.

(40) Barth, E.; Kuczera, K.; Leimkuhler, B.; Skeel, R. D. Algorithms for Constrained Molecular Dynamics. J. Comput. Chem. 1995, 16 (10), 1192–1209. 10.1002/jcc.540161003.

(41) Hopkins, C. W.; Le Grand, S.; Walker, R. C.; Roitberg, A. E. Long-Time-Step Molecular Dynamics through Hydrogen Mass Repartitioning. J. Chem. Theory Comput. 2015, 11 (4), 1864–1874. 10.1021/ct5010406.

(42) Eastman, P.; Swails, J.; Chodera, J. D.; McGibbon, R. T.; Zhao, Y.; Beauchamp, K. A.; Wang, L.-P.; Simmonett, A. C.; Harrigan, M. P.; Stern, C. D.; Wiewiora, R. P.; Brooks, B. R.; Pande, V. S. OpenMM 7: Rapid Development of High Performance Algorithms for Molecular Dynamics. PLoS Comput. Biol. 2017, 13 (7), e1005659. 10.1371/journal.pcbi.1005659.

(43) Onufriev, A.; Bashford, D.; Case, D. A. Exploring Protein Native States and Large-scale Conformational Changes with a Modified Generalized Born Model. Proteins: Structure, Function, and Bioinformatics 2004, 55 (2), 383–394. 10.1002/prot.20033.

(44) Gaynes, R. The Discovery of Penicillin—New Insights After More Than 75 Years of Clinical Use. Emerg. Infect. Dis. 2017, 23 (5), 849–853. 10.3201/eid2305.161556.

(45) Langan, P. S.; Vandavasi, V. G.; Weiss, K. L.; Cooper, J. B.; Ginell, S. L.; Coates, L. The Structure of Toho1 Β-lactamase in Complex with Penicillin Reveals the Role of Tyr105 in Substrate Recognition. FEBS Open Bio 2016, 6 (12), 1170–1177. 10.1002/2211-5463.12132.

(46) Zain, R.; Vihinen, M. Structure-Function Relationships of Covalent and Non-Covalent BTK Inhibitors. Front. Immunol. 2021, 12. 10.3389/fimmu.2021.694853.

(47) Bender, A. T.; Gardberg, A.; Pereira, A.; Johnson, T.; Wu, Y.; Grenningloh, R.; Head, J.; Morandi, F.; Haselmayer, P.; Liu-Bujalski, L. Ability of Bruton’s Tyrosine Kinase Inhibitors to Sequester Y551 and Prevent Phosphorylation Determines Potency for Inhibition of Fc Receptor but Not B-Cell Receptor Signaling. Mol. Pharmacol. 2017, 91 (3), 208–219. 10.1124/mol.116.107037.

(48) Stille, J. K.; Tjutrins, J.; Wang, G.; Venegas, F. A.; Hennecker, C.; Rueda, A. M.; Sharon, I.; Blaine, N.; Miron, C. E.; Pinus, S.; Labarre, A.; Plescia, J.; Burai Patrascu, M.; Zhang, X.; Wahba, A. S.; Vlaho, D.; Huot, M. J.; Schmeing, T. M.; Mittermaier, A. K.; Moitessier, N. Design, Synthesis and in Vitro Evaluation of Novel SARS-CoV-2 3CLpro Covalent Inhibitors. Eur. J. Med. Chem. 2022, 229, 114046. 10.1016/j.ejmech.2021.114046.

(49) Pillaiyar, T.; Manickam, M.; Namasivayam, V.; Hayashi, Y.; Jung, S.-H. An Overview of Severe Acute Respiratory Syndrome–Coronavirus (SARS-CoV) 3CL Protease Inhibitors: Peptidomimetics and Small Molecule Chemotherapy. J. Med. Chem. 2016, 59 (14), 6595–6628. 10.1021/acs.jmedchem.5b01461.

(50) Lee, T.-W.; Cherney, M. M.; Huitema, C.; Liu, J.; James, K. E.; Powers, J. C.; Eltis, L. D.; James, M. N. G. Crystal Structures of the Main Peptidase from the SARS Coronavirus Inhibited by a Substrate-like Aza-Peptide Epoxide. J. Mol. Biol. 2005, 353 (5), 1137–1151. 10.1016/j.jmb.2005.09.004.

(51) Martina, E.; Stiefl, N.; Degel, B.; Schulz, F.; Breuning, A.; Schiller, M.; Vicik, R.; Baumann, K.; Ziebuhr, J.; Schirmeister, T. Screening of Electrophilic Compounds Yields an Aziridinyl Peptide as New Active-Site Directed SARS-CoV Main Protease Inhibitor. Bioorg. Med. Chem. Lett. 2005, 15 (24), 5365–5369. 10.1016/j.bmcl.2005.09.012.

(52) Rarey, M.; Kramer, B.; Lengauer, T.; Klebe, G. A Fast Flexible Docking Method Using an Incremental Construction Algorithm. J. Mol. Biol. 1996, 261 (3), 470–489. 10.1006/jmbi.1996.0477.

(53) Yang, H.; Yang, M.; Ding, Y.; Liu, Y.; Lou, Z.; Zhou, Z.; Sun, L.; Mo, L.; Ye, S.; Pang, H.; Gao, G. F.; Anand, K.; Bartlam, M.; Hilgenfeld, R.; Rao, Z. The Crystal Structures of Severe Acute Respiratory Syndrome Virus Main Protease and Its Complex with an Inhibitor. Proceedings of the National Academy of Sciences 2003, 100 (23), 13190–13195. 10.1073/pnas.1835675100.

(54) Warrensford, L.; Pittman, A. R.; Austin, S.; Woodcock, H. L. CovCIFDock: Covalent Docking with CIFDock and Hybrid QM/MM Minimizations. March 5, 2025. 10.26434/chemrxiv-2025-mcrsz.

(55) Wu, Q.; Huang, S.-Y. HCovDock: An Efficient Docking Method for Modeling Covalent Protein–Ligand Interactions. Brief. Bioinform. 2023, 24 (1). 10.1093/bib/bbac559.

(56) Shamir, Y.; Gabizon, R.; Rogel, A.; Lin, D. Y.; Andreotti, A. H.; London, N. Discovery of Covalent Ligands with AlphaFold3. J. Am. Chem. Soc. 2026, 148 (12), 13043–13054. 10.1021/jacs.5c22222.

(57) Röhrig, U. F.; Goullieux, M.; Bugnon, M.; Zoete, V. Attracting Cavities 2.0: Improving the Flexibility and Robustness for Small-Molecule Docking. J. Chem. Inf. Model. 2023, 63 (12), 3925–3940. 10.1021/acs.jcim.3c00054.

