## Supporting Information for "CHARMM-GUI *Covalent Ligand Docker* as a Web-based Molecular Docking Platform for Covalent Ligands"

**Table S1.** Built-in library of warhead transformations supported by CGUI-CLD.

| Warhead Type | Linked Residue | PDB ID | Ligand ID |
|---|---|---|---|
| 1-Hydroxy-2,6-Dioxa-3-Oxophosphorinone | SER | 1sde | 2PB |
| 1,3-Oxazin-6-one | CYS | 6mgu | JQS |
| Acyl Phosphonate | CYS | 4xbb | 3ZR |
| Acyloxymethyl Carbonyl | CYS | 1nmq | 160 |
| Aldehyde | CYS,LYS,SER | 6m0k,4pjf,5uzw | FJC,30W,ZPR |
| Alkyne | CYS | 6sgd | LD5 |
| Alpha-Cyanovinyl Carbonyl | CYS | 4yhf | 4C9 |
| Alpha-Cyclosulfate | ASP | 5npb | 93Z |
| Alpha-Hydroxy Sulfonic Acid | CYS | 4dcd | K36 |
| Alpha-Pyrone | SER | 1k2i | SN1 |
| Amidine | CYS | 3i4a | LN5 |
| Aminium Ion | ARG | 6b7o | AR6 |
| Aryl Sulfone | CYS,SER | 4onm,2pu4 | N2F |
| Aryloxymethyl Carbonyl | CYS | 1rwo | KB2 |
| Azide | LYS | 2zz6 | 6AZ |
| Aziridine | ASP,CYS,GLU | 5i23,2gkj,6sxt | 66V,ZDR,LXE |
| Beta-Lactam | CYS,SER | 4a52,5kmw | IM2,PNN |
| Beta-Lactone | GLU,SER | 4wv7,4fwg | 3UM,SLA |
| Beta-Sulfanylvinyl Carbonyl | CYS | 5w1y | 9SV |
| Beta-Sulfonylvinyl Carbonyl | CYS | 6kx3 | 8ZO |
| Beta-Sulfonylvinyl Nitrile | CYS | 4onn | BY1 |
| Beta-Sultam | SER | 5tx9 | BSA |
| Borate | SER | 2id8 | 2DB |
| Boronic Acid | GLU,HIS,SER,THR | 6ert,6vim,2i72,5inh | BVZ,PBC,VA1,6C1 |
| Butadienyl Carbonyl | CYS | 3x1i | 66B |
| Carbamate | CYS,SER | 5h8i,1uma | N2H,IN2 |
| Carbamide | CYS | 6azs | C5S |
| Carbodiimide | GLU | 2db4 | DCW |
| Carbonate | SER | 3i2f | DBC |
| Carboxylic Acid | CYS,LYS,SER | 4jap,1htp,1p0i | 14U,OSS,BUA |
| Cyanamide | CYS | 6dud,6b88 | HB4 |
| Diazo Compound | LYS | 3rdh | 3RD |
| Diazomethyl Carbonyl | CYS | 2djf | 1ZB |

| | | | |
|---|---|---|---|
| Disulfanyl-Ester | CYS | 2qnz | DFD |
| Disulfide | CYS | 3orx | 1F8 |
| Enamine | CYS | 4gpt | 51K |
| Epoxide | ASP,CYS,GLU,HIS | 5tng,3ioq,5d6e,5d6f | 7XE,E64,57R,94A |
| Ester | CYS,LYS,SER | 4q95,6eyz,1esb | SHV,C5W,BBL |
| Furan | LYS | 1e7u | KWT |
| Gamma-Lactam | SER | 1hv7 | 616 |
| Gamma-Lactone | CYS,SER | 6pzp,6y6u | P7S,ODZ |
| Haloisocyanide | CYS | 6und | QCV |
| Halomethyl Amidine | CYS | 6dge | GBG |
| Halomethyl Carbonyl | ASP,CYS,GLU | 1zrm,5rgm,2vem | BUA,U1D,BBR |
| Hemiacetal | ASP,CYS,GLU | 4ba0,3khu,3rom | 5GF,UPG |
| Hydrazide | CYS,SER | 1ayu,5zhr | 48Z,KOK |
| Imidazolidinone | SER | 6cl8 | MK7 |
| Imidoyl Halide | CYS | 6ffm | D8N |
| Isothiazolinone | CYS | 2mlm | 2W7 |
| Isothiocyanate | CYS | 4ef9 | 4NF |
| Ketone | CYS,HIS,LYS,SER | 2bdl,1b59,4s2c,1zpz | 4PR,2HA,F6R,BUK |
| Nitrile | CYS,LYS,SER | 6rn6,3eww,2i03 | K9Q,U1P,AXD |
| Nitroarene | CYS | 6hfv | G1T |
| O-Acyl Hydroxamic Acid | LYS,SER | 6ovz,1scn | N9M,BAA |
| Phosphate | CYS,HIS,LYS,SER | 5wi5,4dwq,2q2t,3txt | 0V5,5GP,AMP,DFP |
| Phosphonohalogenate | SER | 4jll | SEF |
| Phosphorothioate | SER | 2xqk | VX |
| Propargyl Carbonyl | CYS,HIS | 6o8i,5zwh | LTJ,9KX |
| Sulfonyl Halide | LYS,SER | 4fi8,3tjm | 0UC,PMS |
| Thiirane | GLU | 3i1u | BTW |
| Thiol | CYS | 6r5l | JT2 |
| Thioster | CYS,SER | 5gk1,3dpm | 2K3,LAS |
| Thiosulfonate | CYS | 1qwz | ETM |
| Vinyl Carbonyl | CYS,HIS | 8pta,5ng1 | CIF,ZPN |
| Vinyl Halide | HIS | 9xia | DFR |
| Vinyl Sulfonyl | CYS,LYS | 1m6d,4hjs | MYP,18J |

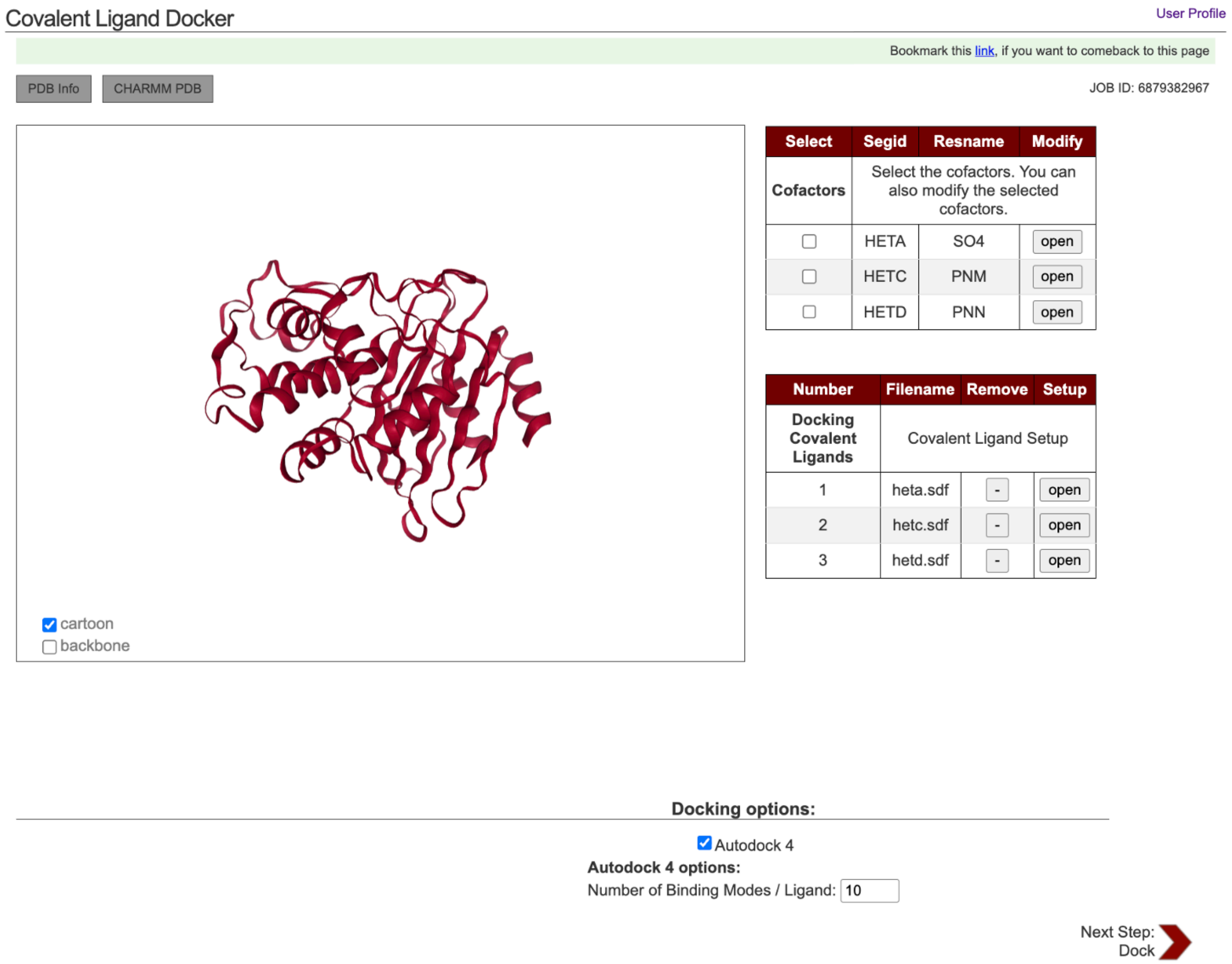


**Figure S1.** CGUI-CLD setup page for covalent ligands and cofactors.

Covalent Ligand Docker

User Profile

Bookmark this link, if you want to comeback to this page

PDB Info | CHARMM PDB

JOB ID: 6879410103

**Docking result:**

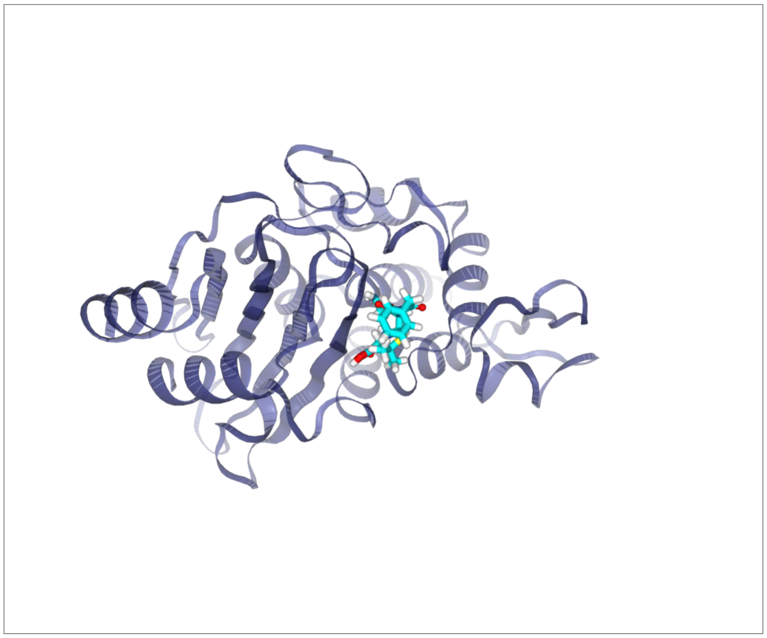

Show All entries

Search:

| View | Package | Ligand | Model | Score | Cluster-Rank | RMSD |
|---|---|---|---|---|---|---|
| ☑ | auto4 | hetc.sdf | 1 | -8.93 | Cluster1-Rank02 | 0.01 |
| ☐ | auto4 | hetc.sdf | 2 | -8.18 | Cluster1-Rank07 | 0.66 |
| ☐ | auto4 | hetc.sdf | 3 | -8.93 | Cluster1-Rank01 | 0.00 |
| ☐ | auto4 | hetc.sdf | 4 | -8.81 | Cluster1-Rank05 | 0.25 |
| ☐ | auto4 | hetc.sdf | 5 | -8.89 | Cluster1-Rank03 | 0.12 |
| ☐ | auto4 | hetc.sdf | 6 | -6.68 | Cluster1-Rank10 | 1.97 |
| ☐ | auto4 | hetc.sdf | 7 | -8.27 | Cluster1-Rank06 | 0.65 |
| ☐ | auto4 | hetc.sdf | 8 | -8.01 | Cluster1-Rank08 | 0.74 |
| ☐ | auto4 | hetc.sdf | 9 | -8.86 | Cluster1-Rank04 | 0.14 |
| ☐ | auto4 | hetc.sdf | 10 | -7.39 | Cluster1-Rank09 | 0.70 |

Showing 1 to 10 of 10 entries

Transfer Selected Complexes to High-Throughput Simulator

**Figure S2.** CGUI-CLD result page with 3D visulization, docking scores, and ranking information.

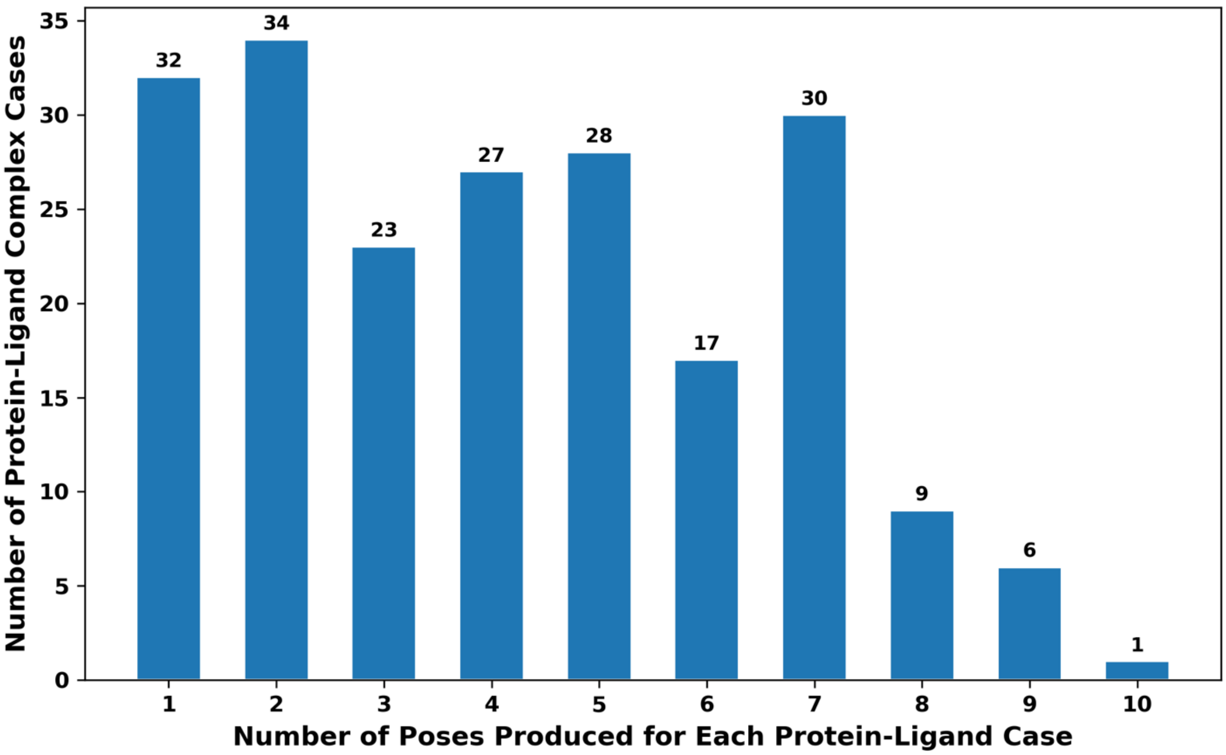


**Figure S3.** Distribution to show how many protein-ligand complex cases produced how many poses.

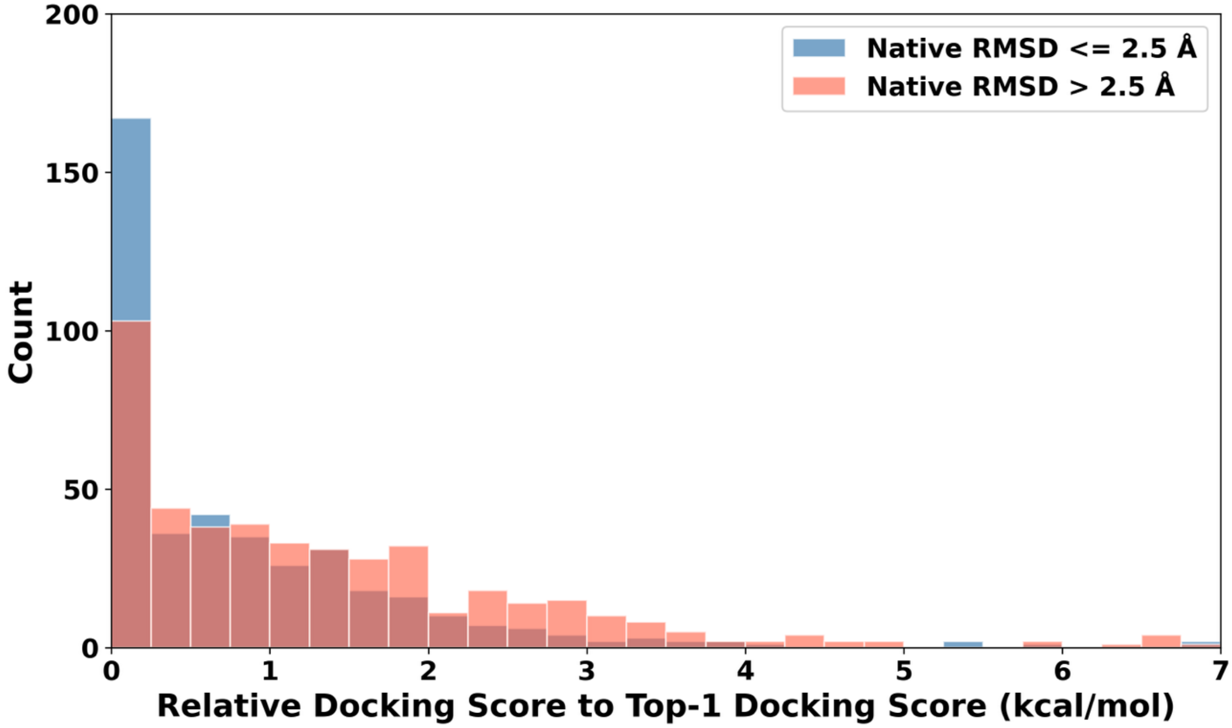


**Figure S4.** Histogram distributions of relative docking score difference to the Top-1 score in each testcase, covering 865 poses. The ligand poses are categorized based on its RMSDs to their native structures (native RMSD): blue for native RMSD <= 2.5 Å (n=414) and red for native RMSD > 2.5 Å (n=451).
